# Tyrosine phosphorylation is critical for ACLY activity in lipid metabolism and cancer

**DOI:** 10.1101/2020.01.20.910752

**Authors:** Johnvesly Basappa, Mahmoud A. ElAzzouny, Delphine C.M. Rolland, Anagh A. Sahasrabuddhe, Kaiyu Ma, Gleb A. Bazilevsky, Steven R. Hwang, Venkatesha Basrur, Kevin P. Conlon, Nathanael G. Bailey, John K. Frederiksen, Santiago Schnell, Yeqiao Zhou, David Cookmeyer, Jan M. Pawlicki, Amit Dipak Amin, James L. Riley, Robert B. Faryabi, Jonathan H Schatz, Kathryn E. Wellen, Ronen Marmorstein, Charles F. Burant, Kojo S.J. Elenitoba-Johnson, Megan S. Lim

## Abstract

A fundamental requirement for growth of rapidly proliferating cells is metabolic adaptation to promote synthesis of biomass^1^. ATP citrate lyase (ACLY) is a critical enzyme responsible for synthesis of cytosolic acetyl-CoA, the key building component for *de novo* fatty acid synthesis and links vital pathways such as carbohydrate and lipid metabolism^2^. The mechanisms of ACLY regulation are not completely understood and the regulation of ACLY function by tyrosine phosphorylation is unknown. Here we show using mass-spectrometry-driven phosphoproteomics and metabolomics that ACLY is phosphorylated and functionally regulated at an evolutionary conserved residue, Y682. Physiologic signals promoting rapid cell growth such as epidermal growth factor stimulation in epithelial cells and T-cell receptor activation in primary human T-cells result in rapid phosphorylation of ACLY at Y682. *In vitro* kinase assays demonstrate that Y682 is directly phosphorylated by multiple tyrosine kinases, including ALK, ROS1, SRC, JAK2 and LTK. Oncogenically activating structural alterations such as gene-fusions, amplification or point mutations of ALK tyrosine kinase result in constitutive phosphorylation of ACLY in diverse forms of primary human cancer such as lung cancer, anaplastic large cell lymphoma (ALCL) and neuroblastoma. Expression of a phosphorylation-defective ACLY-Y682F mutant in NPM-ALK+ ALCL decreases ACLY activity and attenuates lipid synthesis. Metabolomic analyses reveal that ACLY-Y682F expression results in increased β-oxidation of ^13^C-oleic acid-labeled fatty acid with increased labeling of +2-citrate (p<0.01) and +18-oleyol carnitine (p<0.001). Similarly, oxygen consumption rate (OCR) is significantly increased in cells expressing ACLY-Y682F (p<0.001). Moreover, expression of ACLY-Y682F dramatically decreases cell proliferation, impairs clonogenicity and abrogates tumor growth *in vivo*. Our results reveal a novel mechanism for direct ACLY regulation that is subverted by multiple oncogenically-activated tyrosine kinases in diverse human cancers. These findings have significant implications for novel therapies targeting ACLY in cancer and metabolism.

---

ACLY, a key metabolic enzyme involved in glucose and lipid metabolism, connects glycolysis to lipid synthesis and plays an important role in metabolism. The conversion of glucose to fatty acids is dependent on the activity of ACLY which converts mitochondria-derived citrate to cytosolic acetyl-CoA^3^. Acetyl-CoA is an important precursor not only for lipid synthesis but also for cell growth and proliferation by promoting the acetylation of histones for gene transcription^4^. Cell proliferation requires a constant supply of lipids and lipid precursors to fuel membrane biogenesis and protein modification^5^. Glutamine can also be converted to citrate by the reversal of the Krebs cycle catalyzed by isocitrate dehydrogenase and aconitase^6^. ACLY supports *de novo* lipid synthesis and knockdown of ACLY reduces the ability of cells to metabolize glucose to lipids^6^. The role of ACLY in tumor growth has been substantiated by observation that the small molecule inhibitor of ACLY, SB204990, or shRNA mediated knockdown of ACLY abrogate the tumor growth in xenograft models^2^.

In living cells, the activity of ACLY is dependent on its homotetramerization. The complete crystal structure of ACLY has been recently solved by two independent groups^7,8^. We and others have also recently reported cryo-electron microscopy structures of human ACLY, with and without bound acetyl-CoA (Wei X Nat Struct Mol Biol 2019). ACLY activity is also regulated by post-translational modifications including ubiquitination, acetylation^9^ and serine phosphorylation^10^. While a functional role for serine phosphorylation via AKT has been demonstrated to enhance ACLY activity and regulate histone acetylation in human gliomas and prostate tumors^11^, no regulatory role for tyrosine phosphorylation has been established for ACLY function. Receptor tyrosine kinases are key regulators of critical cellular processes such as proliferation, differentiation, cell survival, migration, cell-cycle control and metabolism^12^. Mutational activation of tyrosine kinases resulting in aberrant activation of phosphorylation-mediated intracellular signaling are causally linked to several diseases including cancer, diabetes, inflammation and angiogenesis. Tyrosine kinase (including ALK)-mediated post-translational modification of metabolic enzymes such as pyruvate kinase M2 (PKM2) regulates tumor metabolism^12^. However, a functional role of tyrosine-kinase mediated phosphorylation of ACLY has not been described and the effects on tumor metabolism and growth are unknown.

In our present study, we employed a mass spectrometry-based global phosphoproteomic and metabolomics strategy to elucidate novel mechanisms of oncogenic tyrosine kinase-mediated regulation of metabolic pathways. Our results implicate a direct link between tyrosine kinase-mediated signaling and lipid metabolism that is frequently subverted by oncogenic driver tyrosine kinases in diverse forms of human cancer.

To discover novel mechanisms of phosphotyrosine-mediated oncogenesis, we performed phosphoproteomic analysis of eighteen lymphoma-derived cell lines including SU-DHL-1, SUP-M2 and Karpas 299 harboring the t(2;5)(p23;q35) aberration which encodes the oncogenic chimeric fusion tyrosine kinase NPM-ALK, and generated a compendium of phosphorylated proteins with site mapping of 881 phoshorylated tyrosine peptides. To identify tyrosine phosphorylation events that could be driven by constitutive ALK activation, we performed clustering of pairwise correlations of 359 phosphotyrosine residues in the phosphoproteomic dataset that were measured in at least 3 out of the 18 cell lines. We constructed a heatmap based on Pearson correlation between phosphorylated tyrosine residues that were correlated with phosphorylation of ALK Y1604 (activated ALK) (Extended Data Fig. 1a). We observed that several tyrosine residues, ALK Y1584, ALK Y1507, ALK Y1131, ALK Y1096, PKM Y105, SHC1 Y427 and WDR1 Y238 whose phosphorylation have been reported in ALK positive ALCL^12^ were highly correlated with ALK Y1604 phosphorylation (Extended Data Fig. 1a). Notably ACLY Y131 (Pearson correlation: 0.79, p-value = 0.0001035) and Y682 phosphorylation (Pearson correlation: 0.79, p<1E-4) was significantly correlated with ALK Y1604 (activated ALK) (Extended Data Fig. 1a). To determine whether ACLY Y682 is regulated by ALK activity, we subjected the NPM-ALK positive SU-DHL-1 cells to a selective small molecule inhibitor of ALK (CEP-26939) followed by global phosphoproteomic analysis^13,14^. We observed reduction of phosphorylation of several ALK tyrosine residues including pY1604, pY1078, pY1092, pY1096, pY1131, pY1507, pY1584 (Extended Data Fig. 1c) and ALK-regulated phosphotyrosine substrates including PKM2^15,16^ WDR1^16^ and SHC1^17^ (Extended Data Fig. 1b). Importantly, we observed significant reduction (p <0.05) of ACLY Y682 after ALK inhibition compared to DMSO control (Extended Data Fig. 1b). Global phosphoproteomic analysis using another ALK inhibitor (Crizotinib) corroborated the observations with CEP-26939 (Extended Data Fig. 2).

**Figure 1:**
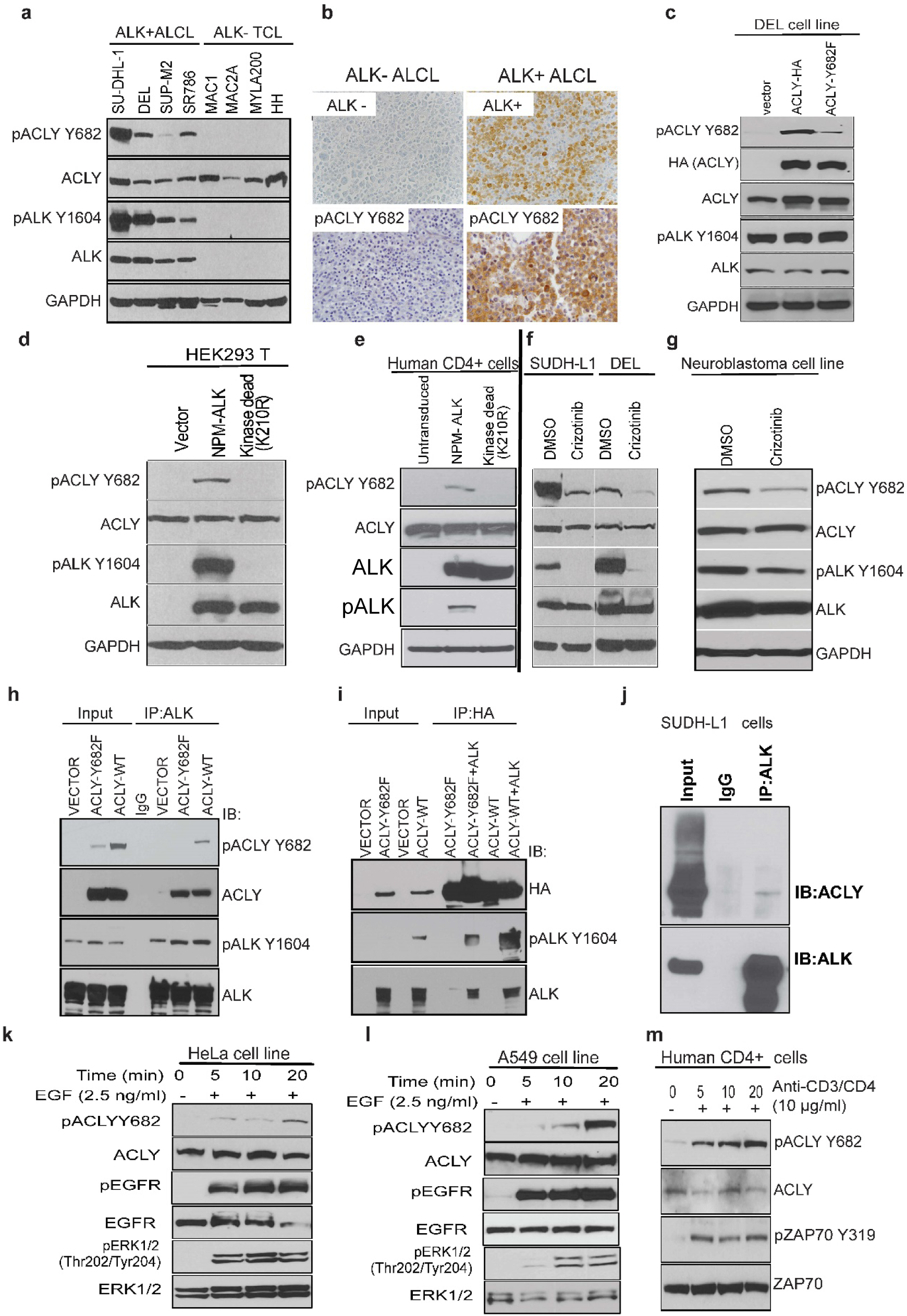
NPM-ALK interacts with ACLY and phosphorylates on Y682 residue in ALK+ALCLs. a, Western blot analyses of pACLY, ACLY in ALK+ALCL and ALK-negative T-cell lymphoma. b, Immunohistochemical expression of pACLY Y682 (lower panel) in human primary ALK+ALCL (right panel) and ALK-T cell lymphoma biopsy samples (left panel). c, DEL stable cell lines expressing vector, ACLY-WT and ACLY-Y682F. d, HEK293T cells transfected with vector, active NPM-ALK and kinase-defective NPM-ALK (K210R). e, Human primary CD4+ cells transduced with active NPM-ALK and kinase-defective NPM-ALK (K210R). f. ALK+ALCL cell lines SU-DHL1 and DEL cells treated with small molecule inhibitor of ALK, crizotinib 300 nM or DMSO for 6 hours. g. ALK+ neuroblastoma cell line NB1 treated with small molecule inhibitor of ALK, crizotinib or DMSO for 6 hours. h. HEK293T cells co-transfected with ACLY-WT, ACLY-Y682F alone with active NPM-ALK and lystaes were immunoprecipitated (IP) with ALK antibody. i, Reciprocal IP with anti-HA from the previous samples and probed with indicated antibodies. j, SU-DHL1 cell lysates were immunoprecipitated with ALK antibody and probed with ACLY. k,l, Stimulation of EGFR positive cells with EGF induces ACLY Y682 phosphorylation in HeLa cells and A549 cells. m, Stimulation of human primary T-cells with anti-CD3/CD4 increases ACLY Y682 phosphorylation

**Figure 2:**
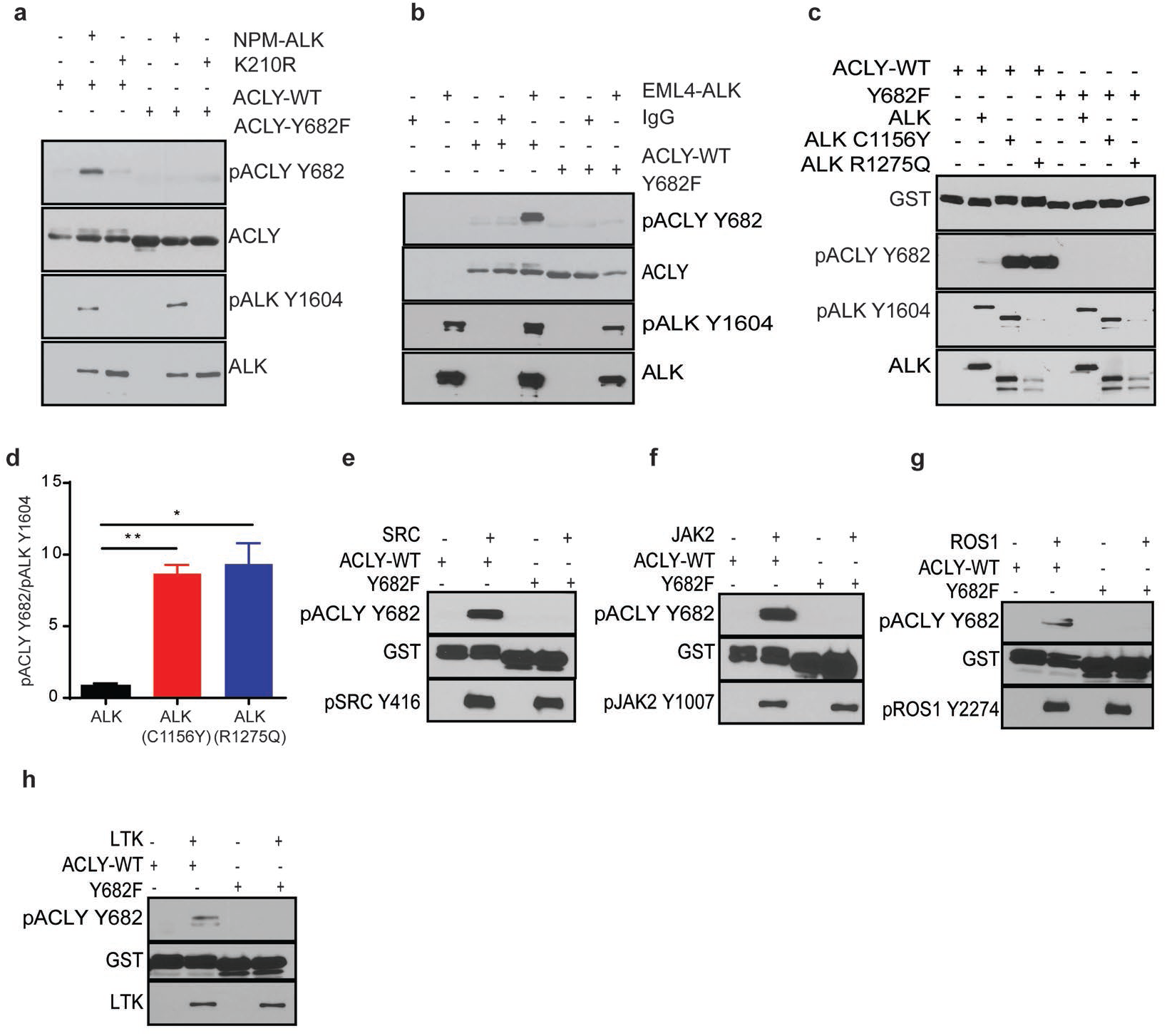
Oncogenic ALK and other tyrosine kinases directly phosphorylate ACLY Y682. a, *In vitro kinase* assay using immunoprecipitated NPM-ALK from ALK+ALCL cell lines and purified ACLY-WT and Y682F proteins. b, *In vitro kinase* assay using immunoprecipitated EML4-ALK from ALK+ non-small cell lung cancer (NSCLC) cell lines and purified ACLY-WT and Y682F proteins. c, *In vitro kinase* assay using purified wild type ALK, ALK C1156Y (mutation in EML4-ALK) and ALK R1275Q (mutation in neuroblastoma). d, *In vitro kinase* assay using purified wild type ALK, ALK C1156Y (mutation in EML4-ALK) and ALK R1275Q (mutation in neuroblastoma). The bar graph represents the relative intensity of the specific bands of pACLY Y682, which are normalized to the value of pALK Y1604 level. e, *In vitro kinase* assay using SRC kinase, f, JAK2 kinase, g, ROS1 kinase and h, LTK kinase. Error bars represent mean values ± standard deviation from three independent experiments (*, P < 0.05, **, P < 0.005).

Correspondingly, co-transfection of HA-tagged ACLY with either active NPM-ALK or the kinase-defective NPM-ALK-K210R mutant in HEK293T cells, and anti-HA immunoprecipitates separated on SDS-PAGE (Extended Fig. 1d) followed by mass spectrometry analysis revealed phosphorylation of ACLY Y131 and Y682 only in the presence of active NPM-ALK (Extended Data Fig. 1e). Cross-species sequence alignment indicated that ACLY Y682 is highly conserved from humans to *C. elegans* (Extended Data Fig. 1f). These results led us to hypothesize that ACLY is novel substrate of ALK tyrosine kinase. To assess the preferential site of NPM-ALK-mediated phosphorylation of ACLY, we generated HA-tagged ACLY-WT, ACLY-Y131F and ACLY-Y682F mutants which were expressed alone, with active NPM-ALK or kinase-defective NPM-ALK-K210R in HEK293T cells. Western blotting of the ACLY immunocomplex with an anti-phosphotyrosine antibody revealed that NPM-ALK preferentially phosphorylates ACLY at Y682 in comparison to ACLY Y131 residue (Extended Data Fig. 1g). These results were corroborated by direct *in vitro* kinase assays (Extended Data Fig. 1h). Taken together, these studies indicate that NPM-ALK directly phosphorylates ACLY Y682. Therefore, we focused our study on exploring the role of ACLY Y682 phosphorylation and its impact on tumor metabolism.

To further explore the effect of crizotinib on ACLY Y682 phosphorylation, 2 ALK+ ALCL cell lines (SU-DHL-1, DEL) and a neuroblastoma cell line (NB1) were treated with crizotinib and lysates were subjected to immunoblot. Western blotting of the lysates with an anti-pACLY Y682 antibody generated to interrogate its phosphorylation status revealed decreased phosphorylation of ACLY at Y682 with crizotinib treatment in comparison to DMSO control (Fig. f,g). Furthermore, the anti-pACLY Y682 antibody was used to assess the level of phosphorylated ACLY in neoplastic and physiologic conditions. Total protein lysates derived from ALK+ ALCL and ALK- T-cell lymphoma cell lines were immunoblotted with anti-pACLY Y682 which revealed constitutive phosphorylation of ACLY at Y682 in ALK+ ALCLs, but not in ALK-T-cell lymphomas (Fig. 1a). Immunohistochemistry (IHC) performed on human tissue biopsies derived from ALK+ ALCL (n=20) and ALK-ALCL (n=28) patients using pACLY Y682 and ALK antibodies (Fig. 1b) revealed that (19/20) 95% of ALK+ ALCLs expressed pACLY-Y682, while (6/28) 21.42% of ALK-ALCL expressed pACLY Y682 demonstrating significant correlation between phosphorylation of ACLY at Y682 and ALK expression (X^2^=31.35; p-value < 0.01).

Next, we generated an ALK+ ALCL cell line (DEL) with lentivirus-mediated stable expression of empty vector, HA-tagged ACLY-WT or ACLY-Y682F. Immunoblotting with pACLY-Y682 antibody revealed that phosphorylation of ACLY Y682 was increased in DEL cells stably expressing ACLY-WT, while cells expressing ACLY-Y682F exhibited basal ACLY Y682 phosphorylation which was comparable to vector (Fig. 1c). Additionally, we stably transduced active NPM-ALK and kinase defective NPM-ALK-K210R constructs in human primary T cells and lysates were prepared 8 days after transduction. Immunoblotting with pACLY-Y682 antibody revealed phosphorylation of ACLY Y682 only in active NPM-ALK transduced human primary T cells (Fig. 1e).

To evaluate the impact of ACLY Y682 phosphorylation on its enzymatic activity, we employed a malate dehydrogenase-coupled assay^18^ in which protein lysates were used as a source of ACLY. Assessment of enzymatic activity revealed significantly decreased ACLY activity in cells expressing ACLY-Y682F in comparison to ACLY-WT (Fig.3c). Pharmacologic inhibition of ALK using crizotinib at 300 nM for 6 hours in 2 ALK+ ALCL cell lines, SU-DHL-1 and DEL resulted in reduction of ACLY activity by >20% and >25%, respectively (p <0.05) (Fig. 3a,b). Taken together, these results indicate that Y682 phosphorylation is critical for ALK-mediated ACLY activity.

**Figure 3:**
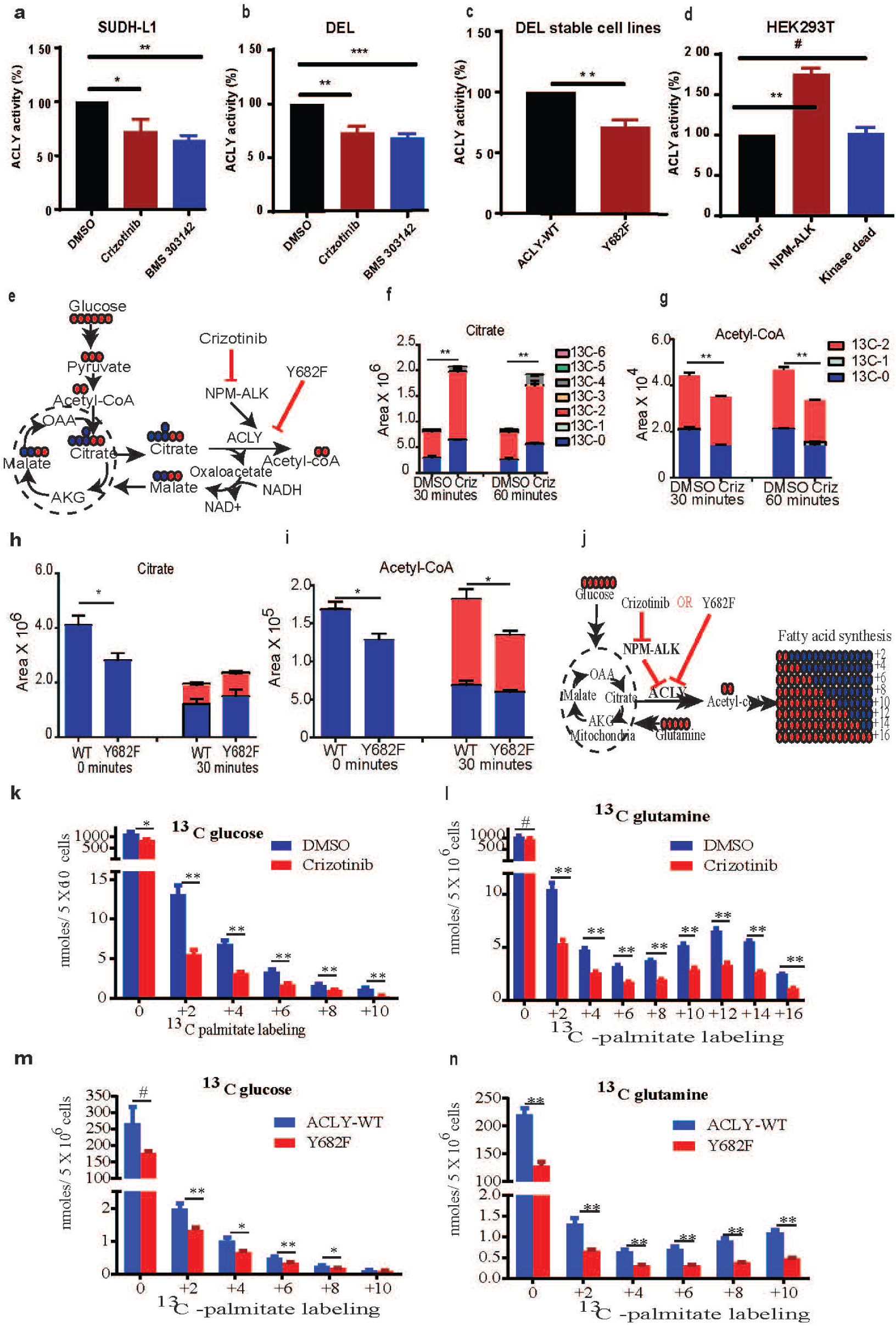
Tyrosine phosphorylation of ACLY increases its activity and lipid metabolism. a, ACLY activity assay on SU-DHL1 cells treated with ALK inhibitor crizotinib and ACLY inhibitor BMS 303142. b, ACLY activity assay on DEL cells treated with ALK inhibitor crizotinib and ACLY inhibitor BMS 303142. c, ACLY activity assay on DEL cells stably expressed ACLY-WT and ACLY-Y682F constructs. d, ACLY activity assay on HEK293T cells transfected with active NPM-ALK and inactive NPM-ALK. e, Schematic representations depicting the flux analysis of ^13^C-glucose. f, ^13^C incorporation in glucose-derived metabolite of citrate in ALK+ALCL cell line (DEL). g, ^13^C incorporation in glucose-derived metabolite of acetyl-CoA. h, ^13^C incorporation in glucose-derived metabolite of citrate in DEL cells stably expressing ACLY-WT and ACLY-Y682F. ^13^C enrichment in glucose-derived metabolite of citrate. i, ^13^C enrichment in glucose-derived metabolite of acetyl-CoA. j, Schematic depicting the flux analysis of ^13^C-glucose and number of carbons labeled (red circles) in fatty acid synthesis. k, ^13^C enrichment in ^13^C-glucose-derived ^13^C -palmitate labeling in DMSO and crizotinib treated ALK+ALCL cell line. l, ^13^C enrichment in ^13^C-glutamine-derived ^13^C -palmitate labeling in DMSO and crizotinib treated ALK+ALCL cell line. m, ^13^C enrichment in ^13^C-glucose-derived ^13^C -palmitate labeling in ACLY-WT and ACLY-Y682F stably transduced DEL, ALK+ALCL cell line. n, ^13^C enrichment in ^13^C-glutamine-derived ^13^C -palmitate labeling in ACLY-WT and ACLY-Y682F stably transduced DEL, ALK+ALCL cell line. Mean ± SEM of triplicates (*, P < 0.05, **, P < 0.005).

To further explore the effect of active NPM-ALK on endogenous ACLY Y682 phosphorylation and its enzymatic activity, we transfected HEK293T cells with empty vector, active NPM-ALK and kinase-defective NPM-ALK-K210R constructs. Western blotting revealed that active NPM-ALK phosphorylated ACLY at Y682 but the kinase-defective NPM-ALK-K210R did not (Fig. 1d). Further, the enzymatic activity of ACLY was dependent on the kinase activity of NPM-ALK. Indeed, cells expressing kinase-defective NPM-ALK-K210R exhibited basal levels of ACLY activity comparable to that seen in empty vector-transfected cells (Fig. 3d). Correspondingly, we further explored the effect of active NPM-ALK on endogenous ACLY Y682 phosphorylation in human CD4+ cells with empty vector, active NPM-ALK and kinase-defective NPM-ALK-K210R constructs and observed similar results (Fig.1e).

To evaluate whether NPM-ALK directly phosphorylates ACLY, we expressed recombinant GST-tagged ACLY-WT and ACLY-Y682F peptides which were co-incubated with active NPM-ALK or kinase-defective NPM-ALK-K210R immunopurified from HEK293T cells that were transfected with respective constructs in *in vitro* kinase assay conditions. Immunoblotting using anti-pACLY Y682-specific antibody revealed that ACLY is phosphorylated at Y682 by NPM-ALK but not by the kinase-defective NPM-ALK -K210R) (Fig. 2a).

To evaluate whether NPM-ALK interacts with ACLY, we transiently transfected vectors encoding HA-tagged ACLY-WT and ACLY-Y682F alone or with active NPM-ALK in HEK293T cells. Immunoblotting of ALK-immunoprecipitation (IP) with anti-HA demonstrated that ACLY-WT and ACLY-Y682F interact with NPM-ALK (Fig. 1g). Further, reciprocal IP with HA revealed NPM-ALK interaction with ACLY-WT and ACLY-Y682F (Fig. 1h). To further evaluate whether endogenous NPM-ALK interacts with endogenous ACLY, we subjected SU-DHL-1 cell lysates for ALK-immunoprecipitation (IP) with ALK antibody and IgG controls and probed with anti-ACLY antibody. These data indicate that ACLY interacts with NPM-ALK in endogenous conditions (Fig.1j).

The gene encoding ALK tyrosine kinase is targeted by multiple oncogenic alterations including translocations/gene fusions, gene amplifications and point mutations that lead to its constitutive activation in diverse human cancers^19^. To determine whether other ALK fusion proteins in addition to NPM-ALK constitutively phosphorylate ACLY Y682, we performed western blot analysis of a non-small-cell lung cancer-derived cell line H3122^20^ that expresses the echinoderm microtubule-associated protein-like 4-anaplastic lymphoma kinase (EML4-ALK) fusion protein. The results revealed that EML4-ALK phosphorylates ACLY Y682 in H3122 cell line (Extended Data Fig. 3a). We then performed *in vitro* kinase assays to evaluate whether EML4-ALK directly phosphorylates ACLY. To this end, we incubated recombinant GST-tagged ACLY-WT and ACLY-Y682F peptides with immunopurified EML4-ALK from HEK293T cells transfected with *EML4-ALK* followed by immunoblotting with anti-pACLY Y682. The results revealed that EML4-ALK directly phosphorylates ACLY Y682 (Fig. 2b). Immunohistochemistry (IHC) using tissue microarrays (TMA) revealed positive reactivity for pACLY Y682 in *EML4-ALK+* primary lung cancer (n=5) (Extended Data Fig. 3b). Furthermore, we detected constitutive phosphorylation of ACLY Y682 in the neuroblastoma cell line (NB1)^21^ (Fig. 1g) which is characterized by *ALK* amplifications.

To investigate whether point mutations in full-length ALK also promote ACLY Y682 phosphorylation, we used recombinant ALK R1275Q which is observed in sporadic and familial neuroblastoma^22^ and ALK C1156Y a known crizotinib resistance mutation in EML4-ALK positive lung cancer^23^. *In vitro* kinase assays revealed increased Y682 phosphorylation in GST-ACLY in the presence ALK R1275Q and ALK C1156Y when compared to wild type full-length ALK (Fig. 2c). In this regard, we quantified the densitometry values and observed a seven-old increase in ACLY Y682 phosphorylation in the presence of *ALK* mutations (Fig. 2d). These findings indicate that activating point mutations in full length ALK (R1275Q) increase Y682 phosphorylation of ACLY and that the crizotinib resistance mutation (C1156Y) led to enhanced ALK-mediated phosphorylation of ACLY Y682. The results indicate that diverse genomic mechanisms of oncogenic ALK activation lead to constitutive phosphorylation of ACLY Y682.

Having observed robust phosphorylation of ACLY Y682 in cells harboring *ALK* gene fusions, amplifications and activating mutations, we sought to investigate whether ACLY could serve as a substrate for other tyrosine kinases. *In vitro* kinase assays using recombinant proteins showed that multiple tyrosine kinases namely SRC (Fig. 2e), JAK2 (Fig. 2f), ROS1 (Fig. 2g) and LTK (Fig. 2h) directly phosphorylate ACLY Y682. Furthermore, IHC demonstrated ACLY Y682 phosphorylation in tissue biopsies of *EGFR* mutant lung cancer (n=5) (Extended Data Fig. 3d) and HER2-amplified breast cancer (n=5) (Extended Data Fig. 3c).

Given our observation of direct and constitutive phosphorylation of ACLY Y682 by multiple oncogenic tyrosine kinases, we reasoned that physiologic cellular growth signals such as growth factor stimulation and T-cell receptor signaling may regulate the tyrosine phosphorylation of ACLY Y682. To test this hypothesis, we stimulated two epithelial cell lines (HeLa and A549) with epidermal growth factor (EGF) and performed western blotting with anti-pACLY Y682. Rapid phosphorylation of ACLY Y682 was observed upon EGF stimulation in a time-dependent fashion in both epithelial cell contexts (Fig. 1,k,l). To further substantiate a role for Y682 phosphorylation in the regulation of physiologic processes, we evaluated pACLY Y682 in primary peripheral blood T cells following stimulation with anti-CD3/CD4^24,25^. These experiments revealed rapid and time-dependent phosphorylation of ACLY Y682 (Fig. 1m). Taken together, these results indicate that diverse physiologic stimuli engaging multiple tyrosine kinase-mediated pathways regulate the phosphorylation of ACLY Y682.

Having established that phosphorylation of ACLY Y682 regulates its enzymatic activity, we sought to evaluate its impact on citrate metabolism. We performed ^13^C-glucose labeling analysis using high-performance liquid chromatography-tandem mass spectrometry (LC-MS/MS) in DEL cells pretreated with DMSO and crizotinib (Schematic shows ^13^C-glucose derived metabolites where carbons are depicted as (red circles) Fig. 3e). As shown in Fig. 2f, ALK inhibition led to time-dependent accumulation of ^13^C_2_-citrate (m+2 red) in comparison to DMSO control. This was associated with concomitant time-dependent reduction of ^13^C_2_-acetyl-Co-A (m+2 red) in response to ALK inhibition by crizotinib (Fig. 2g). Similar results were observed in another ALCL-derived cell line, SUD-HL1 (Extended Data Fig. 4b). This was associated with concomitant time-dependent reduction of ^13^C_2_-acetyl-Co-A (m+2 red) and ^13^C_2_-malonyl-co-A (m+2 red) in response to ALK inhibition (Extended data Fig.5c,d). Similarly, stable expression of the ACLY Y682F mutant in the ALK+ALCL cell line (DEL) led to accumulation of ^13^C_2_-citrate (m+2) and reduced ^13^C_2_-acetyl-Co-A (m+2) compared to ACLY-WT (Fig. 3h,i).

**Figure 4:**
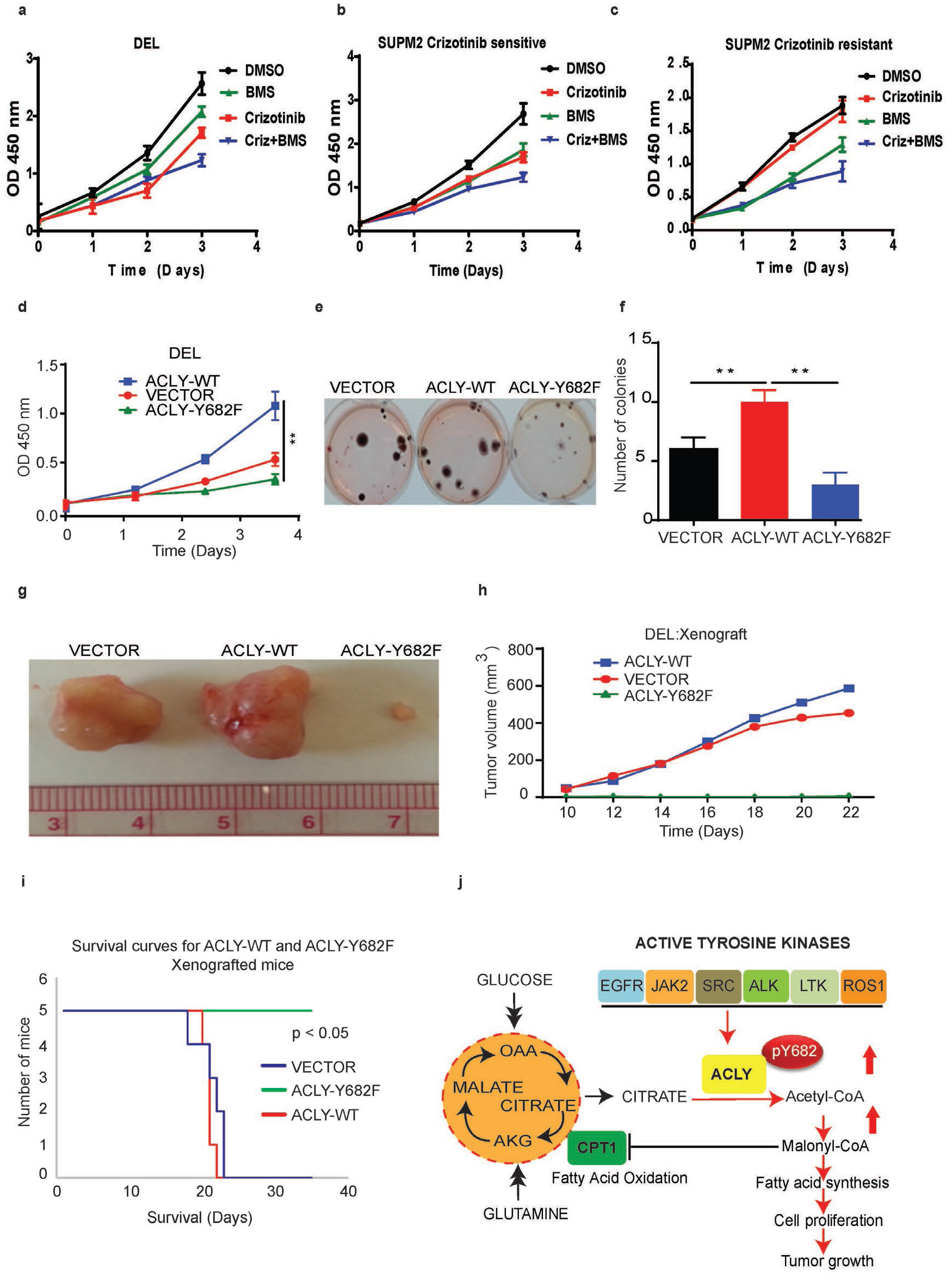
Tyrosine phosphorylation of ACLY regulates cell proliferation and tumor growth. a-c, Cell proliferation of ALK+ALCL cell lines treated with DMSO, ALK inhibitor (crizotinib) and ACLY inhibitor (BMS 303141) using WST-1 assay. a, DEL, b, SUPM2 crizotinib sensitive, c, SUPM2 crizotinib resistant d, Cell proliferation of DEL cells stably expressing vector, ACLY-WT and ACLY-Y682F using WST-1 assay. e, Methylcellulose colony formation assay using DEL cells stably expressing vector, ACLY-WT and ACLY-Y682F. f, Samples analyzed in triplicate, with a representative image (e) and bar graph. g, Representative images of tumors derived from DEL cells stably expressing vector, ACLY-WT and ACLY-Y682F arising from 1X10^7^ cells xenografted into SCID-beige mice. h, Tumor volumes of xenografts derived from DEL cells stably expressing vector, ACLY-WT and ACLY-Y682F in SCID-beige mice. i, Kaplan-Meier survival curves of ALK+ ALCL (DEL) xenografts in SCID-BEIGE mice. j, Schematic summary: Multiple tyrosine kinases phosphorylate ACLY Y682 to regulate its enzymatic activity and promote lipid metabolism and tumor growth.

We next explored the functional role of ALK-mediated ACLY Y682 phosphorylation in *de novo* lipogenesis using metabolomic profiling. DEL cells were cultured in the presence of ^13^C-glucose or ^13^C-glutamine for 24 hours and lipids were extracted and converted into fatty acid methyl esters analyzed by GC-MS (Fig. 3j). These analyses demonstrated that ALK enhances the rate of incorporation of ^13^C-glucose and ^13^C-glutamine derived ^13^C -into palmitate in DEL cells (Fig. 3k,l). This was more pronounced for glutamine-derived ^13^C incorporation (+16) into cellular palmitate relative to glucose-derived ^13^C incorporation (+10 (Fig. 3l). These data indicate that NPM-ALK regulates *de novo* lipid synthesis. The effect of ACLY on *de novo* lipogenesis was then evaluated using DEL cells stably expressing ACLY-WT and ACLY-Y682F cells using similar experimental conditions. As shown in Fig. 3m and Fig. 3n, these experiments revealed that ACLY-Y682F led to significant decrease in ^13^C incorporation into cellular palmitate (p < 0.001) in either ^13^C-glucose- or ^13^C-glutamine-cultured conditions. Taken together, these results indicate that ACLY Y682 phosphorylation promotes *de novo* lipogenesis.

To determine whether phosphorylation of ACLY Y682 impacts fatty acid oxidation, DEL cells stably transduced to express ACLY-WT or ACLY-Y682F mutant were cultured in ^13^C -oleic acid-containing media for LC-MS/MS analysis to quantitate the intermediates derived from β-oxidation of fatty acid (Extended Data Fig. 4e). Complete oxidation of ^13^C -oleic acid in mitochondria yields eight acetyl units which are incorporated in the form of citrate (^13^C_2_-citrate) labeled on two carbons^26^. LC-MS/MS analyses demonstrated increased ^13^C_2_-citrate and oleoyl carnitine (m+18) metabolite levels in cells expressing ACLY-Y682F when compared to ACLY-WT (Extended Data Fig. 4f,g). Taken together, these studies indicate that ACLY Y682 phosphorylation decreases fatty acid oxidation.

The ability to adapt oxygen consumption requirements to favor generation of biomass irrespective of ambient oxygen conditions is a critical necessity during rapid physiologic and oncogenic cell growth^27-29^. To evaluate the role of ACLY Y682 phosphorylation on oxygen consumption rates (OCR), we measured the basal OCR in DEL cell lines stably expressing ACLY-WT and ACLY-Y682F^30^ in the presence of palmitate or oleic acid. These studies showed increased basal OCR in ACLY Y682F cells when compared to ACLY-WT cells (Extended Data Fig. 4h). Taken together these data indicate that phosphorylation of ACLY Y682 favors generation of lipid biomass and anabolic metabolism.

We investigated the effect of small molecule inhibitors of ACLY (BMS 303141)^31^ and ALK (crizotinib)^145^ on cell proliferation of ALK+ALCL cell lines DEL, SUPM2 and SUP-CR500 for 24, 48 and 72 hours. Cells treated with the ACLY inhibitor alone or in combination with crizotinib showed significant reduction of cell proliferation when compared to DMSO control, similar to ALK inhibition (Fig. 4a,b,c). Similarly, we investigated the functional role of phosphorylation of ACLY Y682 in DEL cells stably expressing vector, ACLY-WT or ACLY-Y682F and assessed cell proliferation over 24, 48 and 72 hours. Cells expressing ACLY-Y682F showed significant reduction of proliferation (>69%, p <0.005) when compared to those expressing ACLY-WT (Fig. 4d).

We employed methylcellulose-based colony formation assays to assess the clonogenic potential of ALK positive DEL cells expressing ACLY-WT and ACLY-Y682F mutant using vector only-expressing cells as control. These assays revealed significantly higher (>70%, p<0.005) colony formation in ACLY-WT when compared to ACLY-Y682F mutant cells (Fig. 4e,f). To evaluate *in vivo* tumorigenicity, we established xenografts of DEL cells stably expressing vector control (n=5), ACLY-WT (n=5) and ACLY-Y682F (n=5) by subcutaneous injection of 1X10^7^ cells into SCID-BEIGE mice and examined tumor growth over time. After 21 days, all mice xenografted with vector only and ACLY-WT developed large tumors (>550 mm^3^ volume). By contrast, only one out of five mice injected with ACLY-Y682F cells developed a small tumor *in vivo* while the remaining four mice were tumor-free. The tumor sizes (Fig. 4g) and volumes (Fig. 4h) show significantly diminished tumor growth in ACLY-Y682F expressing DEL cells. Accordingly, whereas all mice harboring ACLY-Y682F expressing DEL xenografts were alive beyond 34 days, all animals xenografted with ACLY-WT and vector-only expressing DEL cells did not survive beyond 23 days (Fig. 4i). Taken together, these results indicate that phosphorylation of ACLY Y682 is critical for tumor growth. Fig. 4j represents the working model in which multiple physiologic and oncogenic tyrosine kinases directly phosphorylate ACLY Y682 to promote lipid metabolism and cellular proliferation.

We show here that phosphotyrosine-mediated regulation of ACLY activity plays a critical functional role in the generation of acetyl-CoA and subsequent lipid metabolism that contribute to tumor growth. Given that cell proliferation requires *de novo* fatty acid synthesis, tyrosine kinase-mediated regulation of ACLY activity provides a direct mechanism for connecting growth factors and other cellular signals such as lymphocyte antigen receptors signaling to lipid metabolism in order to support membrane synthesis required for cell growth. In this regard, our studies demonstrate that Y682 phosphorylation of ACLY is regulated by physiologic signals such as EGF and CD3-mediated T-cell activation. These findings suggest that reversible physiological tyrosine phosphorylation at ACLY Y682 may serve as a switch that directly controls the ability of critical cellular events to immediately trigger lipid synthesis required for cell proliferation. Oncogenically-activated tyrosine kinases are among the most common primary drivers of diverse cancers and may subvert this mechanism by constitutive phosphorylation of Y682 ACLY and stimulation of ACLY activity to promote lipid synthesis and tumor proliferation and may contribute to acquired resistance to small molecular inhibitors. These observations indicate that direct inhibition of ACLY Y682 phosphorylation may offer an attractive therapeutic opportunity for diverse cancers.

## METHODS

### Antibody production

ACLY Y682 phospho-specific, rabbit polyclonal antibody was raised against a KLH-coupled peptide (RTTDGVpYEGVAIG) which corresponds to residue Y682 of human ACLY. Antibody was generated and affinity purified by Pierce Protein Research Services, Rockford, IL.

### Cell lines

Five ALK+ ALCL cell lines (SU-DHL1, DEL, Karpas 299, SUP-M2 and SR786), 2 ALK-ALCL cell lines (MAC1, MAC2A), 2 cutaneous T cell lymphoma (CTCL) cell lines (MYLA, HH), and human embryonic kidney epithelial cell line (HEK293T) were maintained at 37°C in RPMI 1640 (Life Technologies) and DMEM supplemented with 10% fetal bovine serum and Penicillin and Streptomycin (1mM), in a humidified atmosphere containing 5% CO_2_, respectively.

### HA-tagged ACLY-WT and ACLY-Y682F overexpression in HEK293T cells

ACLY was transiently transfected in HEK293T cells using Polyjet (SignaGen Lab) with pLenti vector and pLenti-GIII-CMV-Human-ACLY HA-tagged constructs (Abmgood, Burlington, Canada) by following the manufacturer’s guidelines. Empty vector lacking the ACLY sequence was used as control. ALCY-Y682F mutation was generated by Dpn I-mediated site-directed mutagenesis using Phusion High-Fidelity DNA Polymerase using ACLY-HA as a template.

### ACLY-WT and ACLY-Y682F mutant lentivirus transduction in DEL cells

To stably overexpress ACLY-WT and ACLY-Y682F mutant form of human full length constructs in DEL, lentivirus transduction particles were generated using HEK293T packaging cells transfected by vector plasmid constructs such as psPAX2 and pMD2.G containing full length human ACLY-WT and ACLY-Y682F mutant with HA and GFP tags. The virus particles were transduced into DEL cell line and expanded for three passages; GFP-positive cells were subjected to fluorescence-activated cell sorting (FACS) to obtain cells stably expressing ACLY-WT and ACLY-Y682F mutant.

### Immunoblotting analysis

Proteins were extracted using RIPA lysis buffer containing a cocktail of protease and phosphatase inhibitors. For western blotting, 50 μg of total cell proteins were subjected to SDS-PAGE in 10 or 4-20 % NuPAGE gradient gels under reducing conditions and transferred onto a nitrocellulose membrane. The blots were blocked in 5 % skimmed milk in TBST and probed with primary antibodies overnight. The blots were developed using ECL western blotting detection reagent (GE Healthcare).

### Immunoprecipitation and pull down assays

HA-tagged proteins were immunoprecipitated from HEK293T cells co-transfected with ACLY-WT-HA, ACLY-Y682F-HA with NPM-ALK, NPM-ALK-K201R or EML4-ALK cell lysates by incubation with agarose-HA antibody overnight. For ALK immunoprecipitations, cell lysates from foresaid conditions were incubated with anti-ALK antibody (1:200) and protein A/G agarose overnight. Beads were washed in RIPA buffer and analyzed by immunoblotting.

### Phosphopeptide analysis of ACLY by mass spectrometry (MS) experiments

HA-tagged ACLY-WT alone, or with active NPM-ALK or kinase-defective NPM-ALK-K210R was immunoprecipitated with HA antibody and resolved on SDS-PAGE (NuPAGE 4-20%), the gel was stained with G250 and the bands were excised. Phosphotyrosine containing peptides were purified and subjected to tandem mass spectrometry for protein identification.

### Co-immunoprecipitation and immunoblotting

HEK293T cells were grown in 10-cm plates. After 24 hours, cells were transiently transfected with HA-tagged ACLY-WT, ACLY-Y682F, active NPM-ALK kinase-defective NPM-ALK-K210R, construct expressing EML4-ALK [E6A20] and the empty vector (pCDH-puro) using Polyjet transfection reagent (Signagen) as per the manufacturer’s instructions. Plasmids used to express active NPM-ALK and kinase defective NPM-ALK-K210R has been described previously^24^. The constructs expressing EML4-ALK [E6A20] and the empty vector (pCDH-puro) were generously provided by Dr. Robert C. Doebele, University of Colorado, Denver, CO. Forty-four hours post-transfection, cells were lysed in 1000 μl of RIPA lysis buffer/plate by sonication. Lysates were centrifuged (15,000 × g, 10 min, 4°C) and thesupernatant was incubated overnight at 4°C with anti-HA or anti-ALK antibodies. Fifty μl of protein A/G PLUS-agarose (Santa Cruz Biotechnology) were added for 6 h. After four washes with RIPA lysis buffer, the precipitated proteins were heated in 2 X SDS-PAGE sample buffers at 95°C for 6 min and analyzed by western blotting. Membranes were blocked in TBST plus 5% defatted milk followed by overnight incubation at 4°C with primary antibodies as indicated, i.e. anti-HA, anti-pACLY Y682, anti-ALK and anti-pALK Y1604. Incubation with the appropriate secondary HRP-labeled antibody was followed by detection with ECL western blotting substrate (Roche Applied Science).

### *In vitro* kinase assay using NPM-ALK

Immunoprecipitation of ALK was carried out as described above. Sepharose-bound immune complexes in lysis buffer were washed and resuspended in kinase buffer (50 mM Tris.HCl, pH 7.5, 10 mM MgCl2, 1 mM sodium fluoride, 1 mM sodium orthovanadate, 1mM DTT and 200 μM ATP). GST-ACLY-WT and GST-ACLY-Y682F recombinant peptides were expressed in *E.coli*, BL21 strain and purified by using GST-agarose beads and eluted. Purified GST-ACLY and GST-Y682F peptides were incubated with ALK immunocomplex from NPM-ALK and NPM-ALK (K210R) co-transfected cell lystaes in the presence of kinase buffer and 0.5 mM ATP for 30 min at 30°C. Samples were heated at 95°C for 5 min and separated on a 4-20% gel by SDS-PAGE and followed by western blotting and probed with anti-GST, ant-pACLY Y682, anti-ALK and ant-pALK Y1604 antibodies.

### *In vitro* kinase assays using multiple oncogenic tyrosine kinases

Recombinant GST-ACLY-WT and GST-ACLY-Y682F peptides were subjected to *in vitro* kinase assay using various oncogenic tyrosine kinases as described above. In brief, GST-ACLY-WT and GST-ACLY-Y682F beads were incubated with 100 ng of recombinant ALK, ALK C1156Y (mutated in EML4-ALK), ALK R1275Q (mutated in neuroblastoma), SRC kinase, JAK2, ROS1 and LTK for 30 min at 30°C in kinase buffer. Samples were heated at 95°C for 5 min, separated on a 4-20% gel by SDS-PAGE and followed by western blotting using anti-GST, ant-pACLY Y682, anti-ALK and ant-pALK Y1604, anti-pSRC Y416, anti-pJAK2 Y1007, anti-pROS1 Y2274 and anti-LTK antibodies.

### ACLY enzymatic activity assay

ACLY enzyme activity was determined using the malate dehydrogenase (MDH)-coupled method as described earlier with little modification^9^. Briefly, cell lysates were incubated in reaction buffer containing 10 mM potassium citrate, 10 mM MgCl2, 1 mM DTT, 10 U malic dehydrogenase, 0.3 mM CoASH, 0.1 mM NADH in 50 mM Tris (pH 7.5) and the reaction was initiated by adding 0.2mM ATP in a final volume of 100 ul, incubated at 37°C, and NADH oxidation was continuously monitored every 5 min for 60 min using a microplate reader. To measure the effect of a small molecule inhibitor of ALK, DEL and SU-DHL1 cells were pretreated with crizotinib at 300 nM for 6 hours. For the control experiments, lysis buffer was used in place of cell lysates for nonspecific NADH oxidation. The relative ACLY activities were calculated by normalization to the total protein abundance of the extracts in triplicate.

### Immunohistochemistry (IHC) for pY682-ACLY expression

Formalin fixed, paraffin sections were cut at 5 microns and rehydrated to water. Heat induced epitope retrieval was performed with FLEX TRS Low pH Retrieval buffer (6.10) for 20 minutes. After peroxidase blocking, the rabbit anti p-ACLY Y682 antibody was applied at a dilution of 1:250 at room temperature for 60 minutes. The FLEX HRP EnVision System was used for detection. DAB chromagen was then applied for 10 minutes. Slides were counterstained with hematoxylin.

### ^13^C-glucose isotopomer-labeled metabolomics analysis

DEL cells were treated with DMSO or crizotinib (300 nM) for 3 hours in complete RPMI media containing 10% FBS before ^13^C-glucose flux analysis. Cells were washed in PBS and resuspended at 5 × 10^6^ cells in glucose-free RPMI media containing 10% FBS in the presence or absence of crizotinib for each condition (N=3). The cells were supplemented with U-^13^C-glucose at final concentration of 15 mM and incubated for 30 min and 60 min at 37°C incubator. Cell pellets were snap frozen in liquid nitrogen and processed for LC-MS/MS analysis as detailed in the supplementary materials.

### ^13^C-glucose and ^13^C-glutamine labeling lipid synthesis metabolomics analysis

DEL cells were grown in the presence of DMSO or crizotinib (50 nM) for 24 hours in complete RPMI media containing 10% FBS and ^13^C-glucose (25mM) in three biological replicates. After 24 hours, equal amount of cells (5 X 10^6^ cells) from DMSO and crizotinib-treated flasks were spun down and cell pellets were snap frozen in liquid nitrogen. Similarly, DEL cells were grown in the presence of DMSO or crizotinib (50 nM) for 24 hours in glutamine-free RPMI media containing 10% FBS and ^13^C-glutamine (3mM) in three biological replicates. After 24 hours, equal number of cells (5 X 10^6^ cells) from DMSO and crizotinib-treated flasks were spun down and cell pellets were snap frozen in liquid nitrogen and processed for gas chromatography/mass spectrometry (GC/MS) for analysis of lipid synthesis.

### Fatty acid oxidation

DEL cells stably expressing ACLY-WT and ACLY-Y682F (5 X 10^6^ cells/condition) were seeded in serum/glucose free RPMI medium for 3 hours before the experiment. After 3 hours of starvation, cells were incubated in the presence of 400 μM ^13^C -palmitic acid (Sigma) or 400 μM ^13^C -oleic acids (Sigma) with 2.5 mM glucose in RPMI medium for 2 hours. Cells were immediately snap-frozen using liquid nitrogen and kept at −80°C until extraction of metabolites, as described previously^25^.

### Oxygen consumption rate

Oxygen consumption rate (OCR) was measured using a Seahorse XF24 extracellular flux analyzer (Seahorse Bioscience) as described by the manufacturer protocol. Twenty four hours before the experiment, DEL cells stably expressing ACLY-WT and ACLY-Y682F were cultured in complete RPMI medium. On the day of metabolic flux analysis, the culture medium was replaced with 675 µl of unbuffered serum/glucose free RPMI and seeded on Seahorse XF-24 plates at a density of 1 × 10^5^ cells per well and incubated at 37 °C in a non-CO_2_ incubator for 1 hour. All injection reagents were adjusted to pH 7.4. Baseline rates were measured at 37°C before injecting final volume of 1.0 mM glucose and 400 uM palmitate or oleate. After the addition of 1.0 mM glucose and each fatty acid, OCR readings were automatically calculated from five replicates by the Seahorse XF-24 software.

### EGF stimulation of HeLa and A549 cells

HeLa and A549 cells were grown in 10-cm plates in complete DMEM and RPMI media, respectively for 48 hours. At 80% confluency, cells were serum starved for 16 hours. Cells were stimulated with 2.5 ng/ml EGF for 5, 10 and 20 minutes. Cells were then processed for western blotting as described above with antibodies for anti-EGFR, anti-phospho EGFR, anti-ACLY, anti-ERK1/2 and anti-pERK1/2 and anti-pACLY Y682.

### Anti-CD3/CD4 stimulation of human primary T cells

Human primary T cells were purified from whole blood of healthy donors provided by the Human Immunology Core facility at the University of Pennsylvania. For each condition, 6 x10^6^ cells were stimulated with 10ug/ml of anti-CD3/CD4 antibody in PBS at room temperature for 5, 10 and 20 minutes. Cell lysates were processed for western blotting as described above. The blots were probed with primary antibodies as indicated, i.e. anti-ACLY, anti-phospho ACLY Y682, anti-ZAP70 and anti-pZAP70.

### Culturing, stimulation, and transduction of primary human CD4+ Cells

Primary CD4+T cells were acquired from anonymous donors through the University of Pennsylvania’s Human Immunology Core using an institutional review board-approved protocol. Primary human CD4+ T cells expressing NPM-ALK or NPM-ALK-K210R (NPM-ALK-KD) were prepared as previously described^26^. Transduced T cells were diluted to a concentration of 3.0 x 10^5^ cells/mL every 2 to 3 days. The cells were counted using the electrical sensing zone method (Multisizer 3, Beckman Coulter, Indianapolis, IN)

### Preparation of cell lysates from primary human CD4+ T cells

Untransduced CD4+ T cells or CD4+ T cells expressing NPM-ALK or NPM-ALK-K210R were pelleted and re-suspended in 1 mL of Lysis buffer, Protease inhibitor (75 mM NaCl, 25 mM Tris, 2.5 mM EDTA, 5 mM NaF, 1 mM PMSF, 1 mM Na_3_VO_4_, and 1 complete™protease inhibitor cocktail tablet/25 mL buffer, SigmaAldrich, St Louis, MO) at 4°C for 1 hour while rotating. The lysate was then isolated via centrifugation (10 minutes at 13,793 xg) and extraction of the supernatant. Protein concentrations were then quantified using a Pierce™ Coomassie (Bradford) Protein Assay kit (ThermoFisher) according to standard protocol, after which the lysates were stored at −80°C until desired for use in immunoblotting.

### Cell proliferation and colony formation assay

ALK**+**ALCL cell lines, DEL, SU-DHL1 and Karpas 299, DEL cells expressing vector, ACLY-WT and ACLY-Y682F mutant cells were plated at a concentration of 1 × 10^5^ cells/well in 6 well plates. Cells were treated with DMSO, crizotinib and BMS-303141 alone or in combination for 24, 48 and 72 hours, Cell proliferation was assessed by WST-1 assay as per manufacturer’s protocol (Roche Applied Science, Indianapolis, IN, USA). Colony formation assay was performed with MethoCult methylcellulose-based media as per manufacturer’s protocol (Stemcell Technologies, Vancouver, British Columbia, Canada). After 14 days, colonies were stained with iodonitrotetrazolium chloride overnight and counted as described previously^5^.

### Xenograft model

Four-week-old male SCID-BEIGE mice (CB.17 SCID-BEIGE) (Charles River Laboratory, Wilmington, MA) were used For each condition, a total of 1 × 10^7^ DEL cells expressing vector, stable ACLY-WT or 682F cells were suspended in 100 μl of saline containing 50% Matrigel (BD Biosciences, Becton Drive, NJ) and injected subcutaneously into the flanks of mice (n=5 each). Tumor growth was monitored on alternate days until 3 weeks, and tumor volumes were estimated. All procedures involving mice were approved by the University Committee on the Use and Care of Animals (UCUCA) at the University of Michigan and conform to their relevant regulatory standards.

### Protein extraction and digestion for phosphoproteomic analysis

Cells were lysed in buffer containing 9 M urea/20 mM HEPES pH8.0/0.1% SDS and a cocktail of phosphatase inhibitors. Six milligrams of protein were reduced with 4.5 mM DTT and alkylated with 10 mM iodoacetamide, then digested with trypsin overnight at 37°C using an enzyme-to-protein ratio of 1/50 (w/w). Samples were desalted on a C18 cartridge (Sep-Pak plus C18 cartridge, Waters). Each sample was prepared in triplicate.

### Phosphopeptide enrichment

Metal oxide affinity chromatography (MOAC) was performed to enrich phosphorylated peptides and reduce the sample complexity prior to tyrosine-phosphorylated peptide immunopurification (pY-IP). We used titanium dioxide (TiO_2_) microparticles (Titansphere® Phos-TiO, GL Sciences Inc.). Briefly TiO_2_ microparticles were conditioned with the buffer A (80% ACN/0.4% TFA), then equilibrated with the buffer B (75% buffer A/25% lactic acid). Peptides were loaded twice on TiO_2_ microparticles and washed 2 times with buffer B and 3 times with buffer A. Hydrophilic phosphopeptides were eluted with 5% ammonium hydroxide solution and hydrophobic phosphopeptides were eluted with 5% pyrrolidine solution. The equivalent of 5 mg of protein was further enriched for phosphorylated tyrosine peptides by overnight immunoprecipitation (pY-IP) using a cocktail of anti-phosphotyrosine antibodies (4G10, Millipore; PT-66, Sigma;p-Tyr-100, Cell Signaling Technology).

### Mass spectrometry analysis

Ammonium hydroxide and pyrrolidine eluents were dried (SpeedVac) and reconstituted in 25 μl sample loading buffer (0.1% TFA/2% acetonitrile). Eluent from phosphotyrosine immunoprecipitation was dried and reconstituted in 35 μl of the loading buffer. An LTQ Orbitrap XL (ThermoFisher) in-line with a Paradigm MS2 HPLC (Michrom Bioresources) was employed for acquiring high-resolution MS and MS/MS data. Ten microliters of the phospho-enriched peptides were loaded onto a sample trap (Captrap, Bruker-Michrom) in-line with a nano-capillary column (Picofrit, 75 μm i.d.x 15 μm tip, New Objective) packed in-house with 10 cm of MAGIC AQ C18 reverse phase material (Michrom Bioresource). Two different gradient programs, one each for MOAC and phosphotyrosine immunoprecipitation samples, were used for peptide elution. For MOAC samples, a gradient of 5-40% buffer B (95% acetonitrile/1% acetic acid) in 135 min and 5 min wash with 100% buffer B followed by 30 min of re-equilibration with buffer A (2% acetonitrile/1% acetic acid) was used. For phosphotyrosine immunoprecipitation samples, which were a much less complex mixture of peptides, 5-40% gradient with buffer B was achieved in 75 min followed by 5 min wash with buffer B and 30 min re-equilibration. Flow rate was ∼0.3 μl/min. Peptides were directly introduced into the mass spectrometer using a nano-spray source. Orbitrap was set to collect 1 MS scan between 400-2000 *m/z* (resolution of 30,000 @ 400 *m/z*) in orbitrap followed by data dependent CID spectra on top 9 ions in LTQ (normalized collision energy ∼35%). Dynamic exclusion was set to 2 MS/MS acquisitions followed by exclusion of the same precursor ion for 2 min. Maximum ion injection times were set to 300 ms for MS and 100 ms for MS/MS. Automatic Gain Control (AGC) was set to 1xe^6^ for MS and 5000 for MS/MS. Charge state screening was enabled to discard +1 and unassigned charge states. Technical duplicate data for each of the MOAC elutions (ammonium hydroxide and pyrrolidine) and triplicate data for the phosphotyrosine immunoprecipitation samples were acquired.

### Bioinformatics analysis

RAW mass spectrometric data were analyzed in MaxQuant environment (version 1.5.3.30) and employed Andromeda for database search^27^ The MS/MS spectra were matched against the human Uniprot FASTA database downloaded on 04/27/2016. Enzyme specificity was set to trypsin and a maximum of 2 missed cleavages. Carbamidomethylation of cysteine was set as a fixed modification while methionine oxidation, protein N-acetylation and serine/threonine/tyrosine phosphorylation were set as variable modifications. The required false discovery rate (FDR) was set to 1% both for peptide and protein levels. In addition, “match between runs” option with a window of 1.5 minute was allowed. Log2-transformed centered intensities of phosphopeptides were used for further analysis.

### Statistical analysis

Statistical analysis and graphical presentation was performed using GraphPad Prism 4.0.

## Acknowledgments

We thank Kristina Fields, University of Michigan for her excellent IHC work. This work was supported by the University of Michigan Cancer Center Pilot Grant, the Department of Pathology, University of Michigan and R01 CA140806-01 (MSL), R01 DE119249, R01 CA136905 (KSJ-EJ).

## Author contributions

J.V.B., M.S.L. and K.S.J.E-J., conceived the project, designed the experiments, and wrote the manuscript. J.V.B., M.A and C.F.B. designed the metabolomics experiments, M.A performed the mass spectrometry for identification of metabolites, D.C.M.R, V.B and K.C performed phosphoproteomic analyses. S.S. performed *in*-*silico* modeling and molecular docking studies. G.B., S.R.H., Z.N., V.M, A.S, K.M.,and T.V performed molecular biology experiments, L.Z., Y.Z. and R.B.F. performed bioinformatics analyses, N.G.B and J.K.F evaluated IHC, C.F.B provided the critical reagents and resource for metabolomics study, D.C., J.M.P. and J.L.R performed human T cells transduction experiments, A.D.A generated crizotinib resistant ALK+ cell lines, J.H.S., R.M., K.E.W provided the critical insights and reagents, K.E.J and M.L supervised and oversaw the study. All authors discussed the results and commented on the manuscript.

---

**Extended Data Figure 1:**
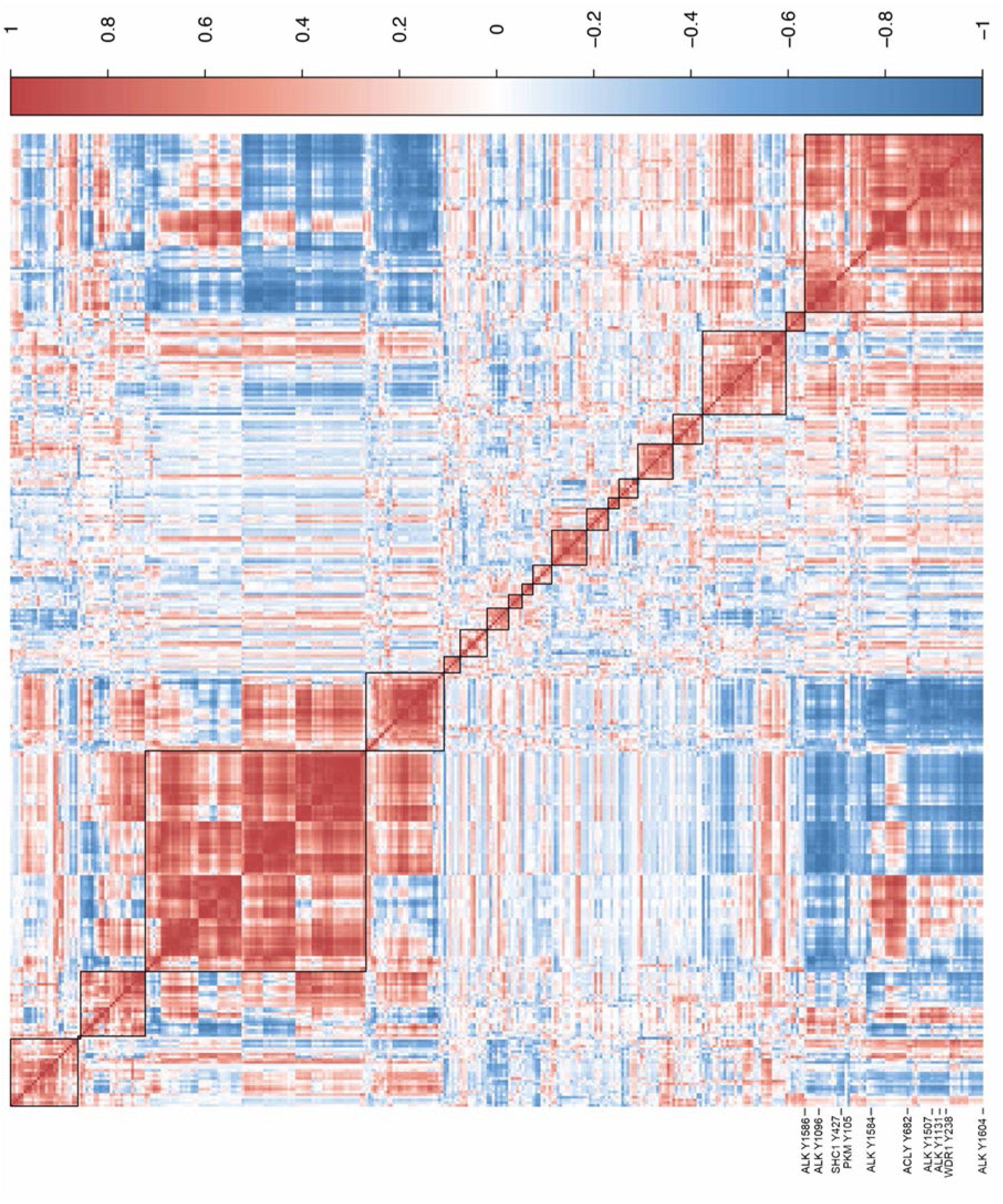
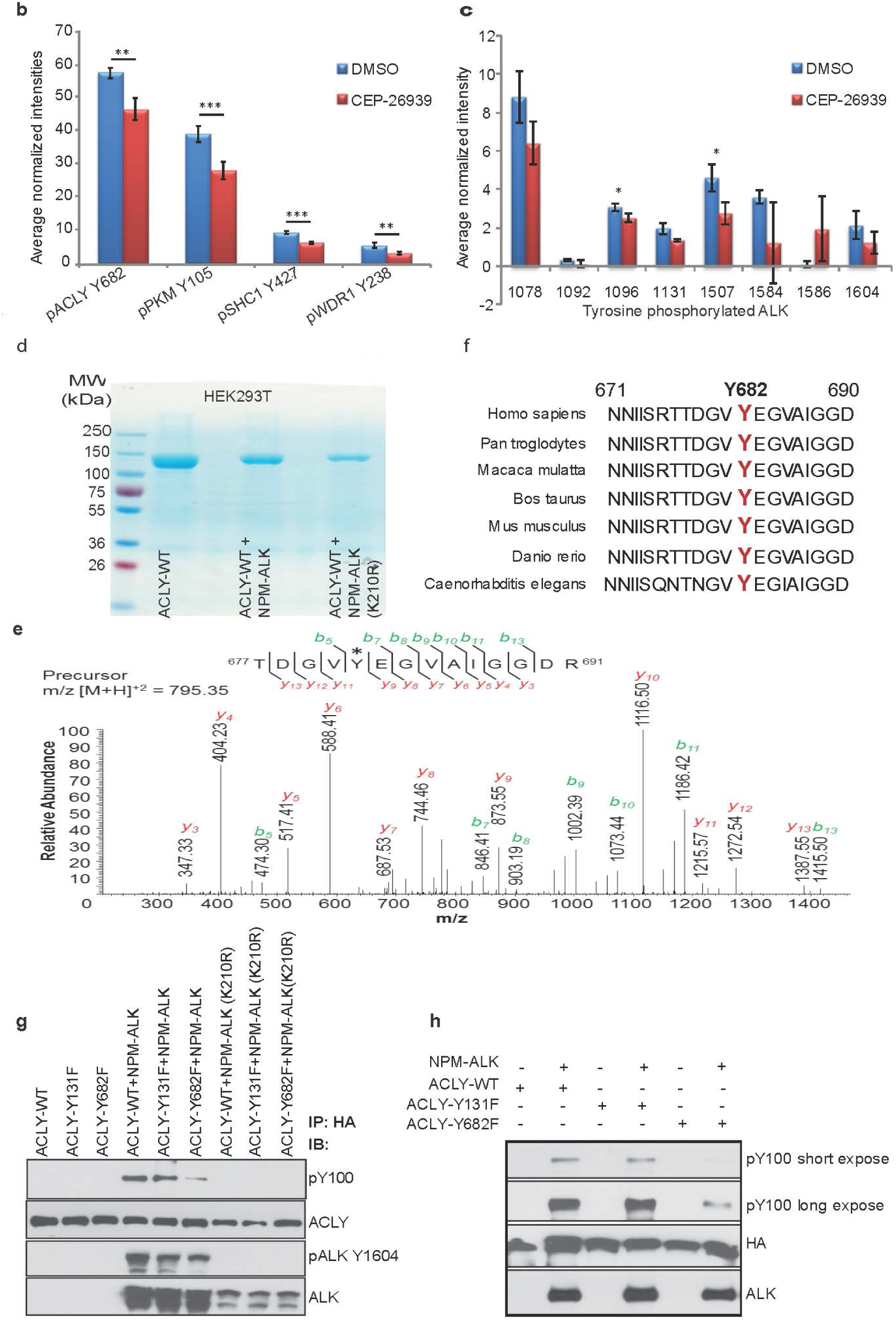
NPM-ALK regulates phosphorylation of ACLY Y682. a, Heat map showing correlation between normalized mass spectral intensity of tyrosine phosphopeptide residues across eighteen lymphoma cell lines. Pearson correlation of the normalized phosphopeptide intensities of 359 tyrosine residues measured in at least 3 cell lines. pACLY Y682 and pALK Y1604 are highly correlated as shown in the bottom right (red) cluster on the heatmap. b, Reduction of pACLY Y682 phosphopeptide intensity in ALK+ cell line (SU-DHL-1) treated with small molecule inhibitor CEP-26939 when compared to DMSO. c, The average normalized intensities of various ALK tyrosine phosphorylated residues in DMSO and CEP-26939 treated SU-DHL-1. ALK+ALCL cell lines (n=3) were processed for phosphopeptide enrichment and analyzed by MS/MS. (p< 0.05 ** and p< 0.01 ***).The bar graph also shows reduced phosphopeptide intensities of known ALK tyrosine kinase substrates such as PKM2, SHC1 and WDR1. (p< 0.05 **, p< 0.01 ***) following ALK inhibition d, HA-tagged ACLY-WT was expressed in 293T cells with active NPM-ALK or kinase-defective NPM-ALK-K210R, immunoprecipitated (IP) with HA and resolved on 4-20% NuPAGE for phosphoproteomic studies. e, Representative MS/MS spectrum of ACLY Y682 phosphorylated peptide revealed by phosphoproteomic analysis. Phosphopeptides isolated through MOAC followed by pY-antibody immnunoprecipitation were resolved on a reverse phase column and collision induced dissociation spectra were obtained using LTQ Orbitrap XL mass spectrometer. A MS/MS spectrum corresponding to ^677^TTDGV**Y**EGVAIGGDR^691^ of ACLY (precursor m/z [M+H]^+2^ = 795.35) is shown. Observed b- and y-ions are indicated. g, Illustration of ACLY Y682 protein sequence homology in multiple species. f, ACLY Y682 site is conserved in all speicies. g, Immunoblotting of ACLY-WT, ACLY-Y131F and ACLY-Y682F mutant constructs expressed in HEK293T cells with indicated antibodies. h, *In vitro* kinase assays of ACLY-WT, ACLY-Y131 and ACLY-Y682F recombinant proteins using endogenous NPM-ALK immunoprecipitated with anti-ALK antibody from ALK+ALCL cell line.

**Extended data Figure 2:**
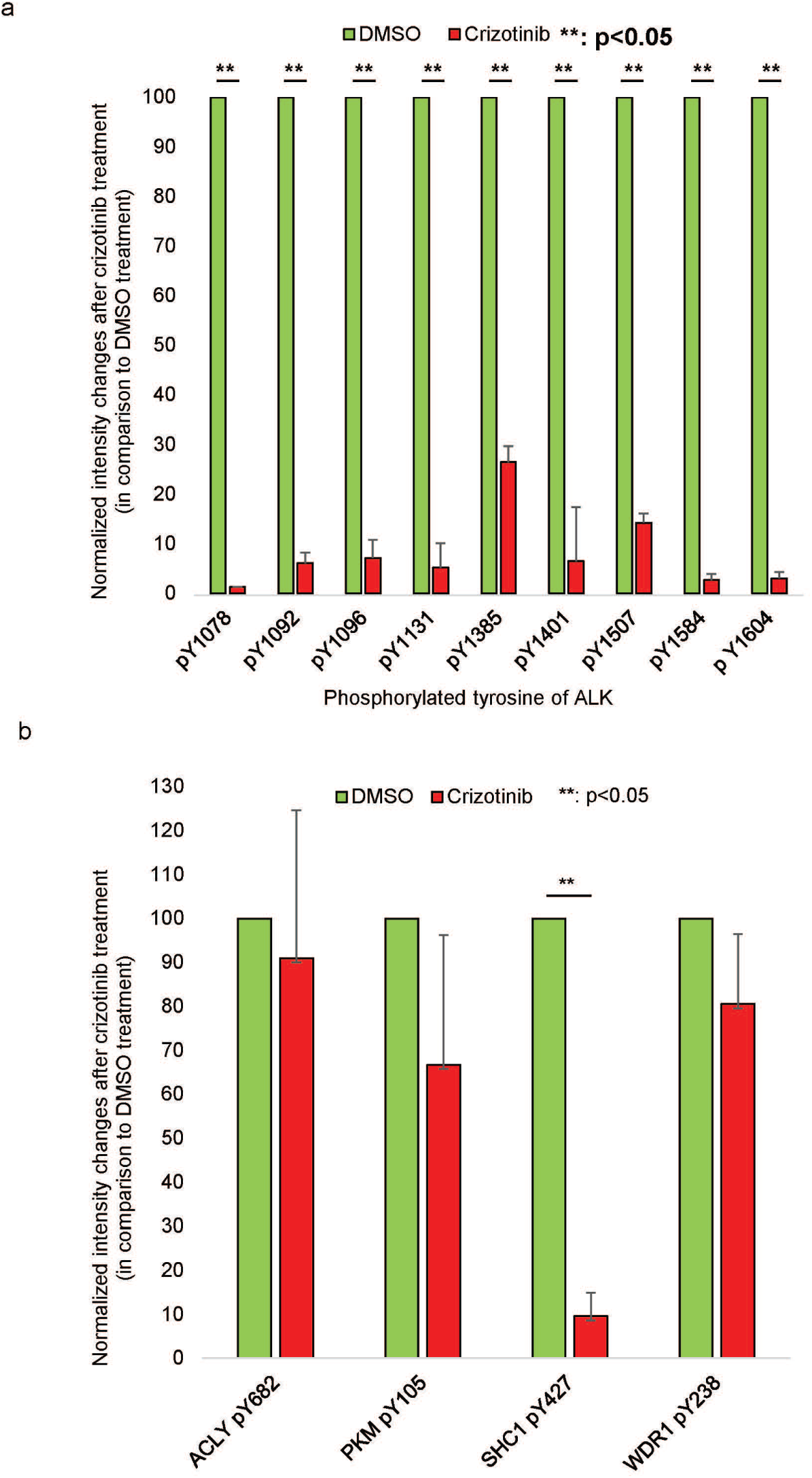
a, Reduction of pACLY Y682 phosphopeptide intensity in ALK+ cell line (SU-DHL-1) treated with small molecule inhibitor crizotinib when compared to DMSO. b, The average normalized intensities of various ALK tyrosine phosphorylated residues in DMSO and crizotinib treated SU-DHL-1. ALK+ALCL cell lines (n=3) were processed for phosphopeptide enrichment and analyzed by MS/MS. (p< 0.05 ** and p< 0.01 ***).The bar graph also shows reduced phosphopeptide intensities of known ALK tyrosine kinase substrates such as PKM2, SHC1 and WDR1. (p< 0.05 **, p< 0.01 ***) following ALK inhibition

**Extended Data Figure 3:**
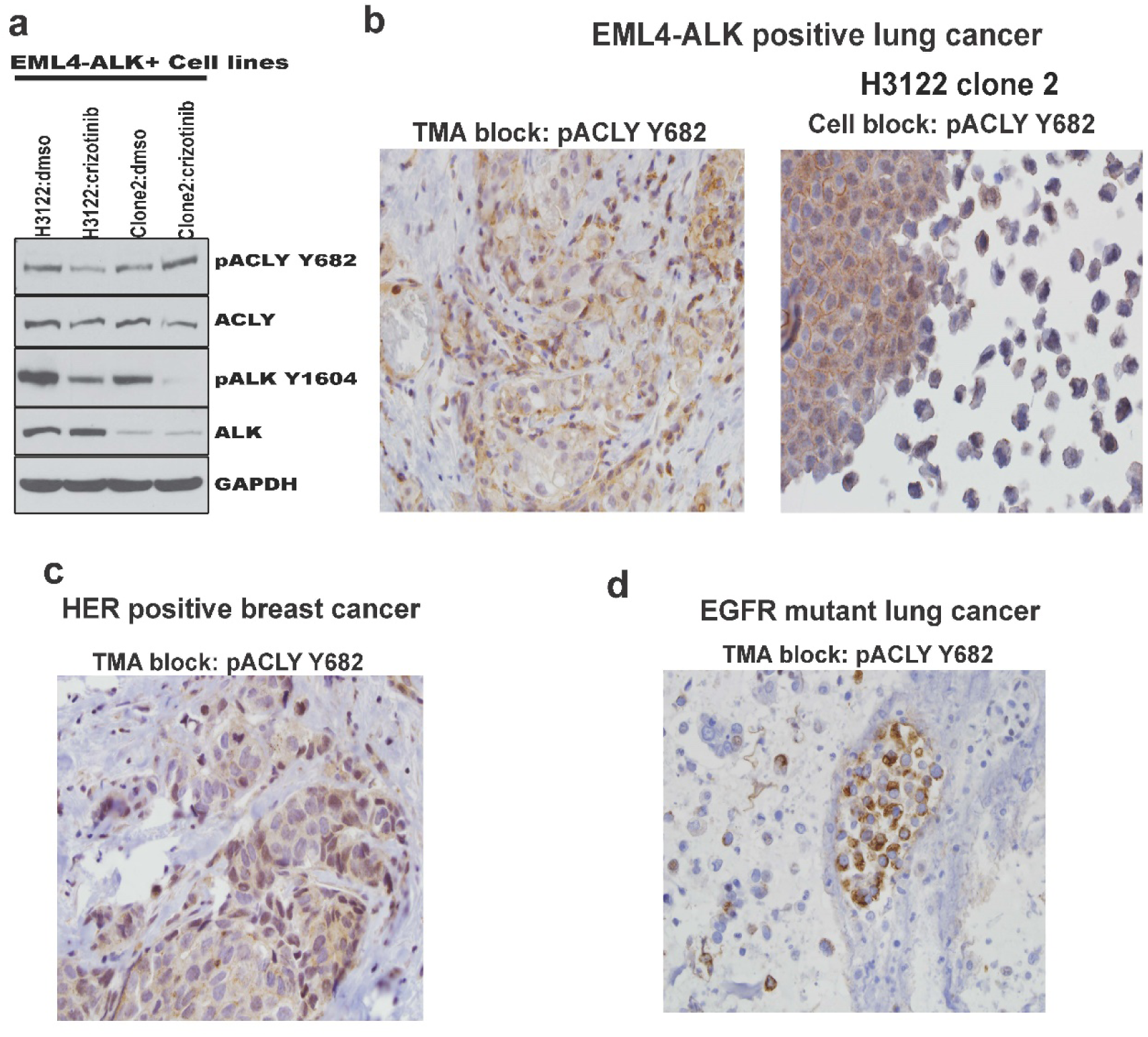
Multiple oncogenic tyrosine kinases regulate ACLY Y682 phosphorylation in human cancer. a, EML4-ALK positive cells (crizotinib H1322 sensitive and clone 2;crizotinib resistant) were treated with DMSO and crizotinib and the lysates were immunoblotted with indicated antibodies. b, Representative image of immunohistochemistry (IHC) of EML4-ALK positive lung cancer biopsy sample (N=5) probed with pACLY Y682. c, Immunohistochemistry of HER-2 amplified breast cancer biopsy probed with pACLY Y682. d, Immunohistochemistry of EGFR mutated lung cancer biopsy probed with pACLY Y682. f, Immunohistochemistry of CD74-ROS1 positive lung cancer biopsy probed with pACLY Y682.

**Extended Data Figure 4:**
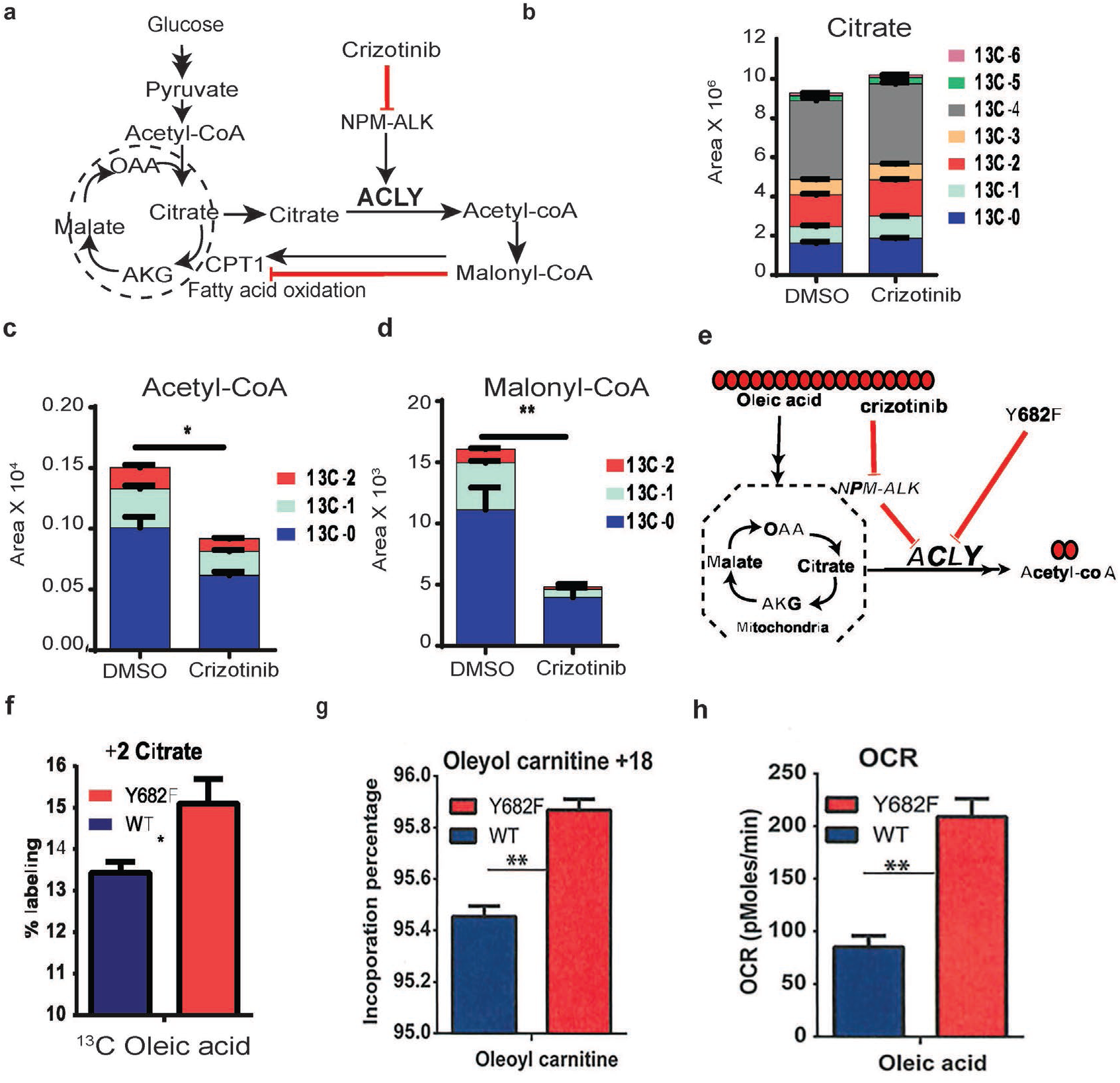
Tyrosine phosphorylation of ACLY attenuates fatty acid oxidation in ALK+ALCL cell lines. a, Schematic representations depicting the flux analysis of ^13^C-glucose in SUD-HL1 cells. b, Increased ^13^C incorporation in glucose-derived metabolite of citrate in crizotinib treated ALK+ALCL cell line (SUD-HL1). c, Decreased ^13^C incorporation in glucose-derived metabolite of acetyl-CoA. d, Decreased ^13^C incorporation in glucose-derived metabolite of malonyl-CoA. e, Schematic representations depicting the flux analysis of ^13^C-oleic acid in DEL cells stably expressing ACLY-WT and ACLY-Y682F constructs. glucose enrichment in glucose-derived metabolite of acetyl-CoA. f, +2 citrate metabolite levels. g, enrichment of oleyol carnitine +18 in ACLY-Y682F cells. h, Measure of oxygen consummation rate (OCR) using seahorse in ACLY-WT and ACLY-Y682F expressing cells. Mean ± SEM of triplicates (*, P < 0.05, **, P < 0.005).

